# Wandering the NPV Maze: Effects of nucleopolyhedrovirus infection on the behavior of Gulf Fritillary (*Dione vanillae*) larvae

**DOI:** 10.64898/2026.06.12.731930

**Authors:** Tiffany Bresnan, Kayla Lizaola, Arietta Fleming-Davies

## Abstract

Parasites can manipulate host behavior to increase their fitness while decreasing host fitness, a phenomenon known as an extended phenotype. Nucleopolyhedroviruses (NPVs), baculoviruses that infect Lepidopteran larvae, have been found to induce vertical climbing behavior and hyperactivity in exposed larvae. We quantified variation in the horizontal wandering behavior induced by different naturally-occurring pathogen isolates in the NPV that infects *Dione* (*Agraulis) vanillae* Linnaeus (Lepidoptera: Nymphalidae). Lab-raised larvae were infected with a constant dose of one of five different field-collected NPV isolates or a water control (n=97 larvae total), and placed in mazes to measure the horizontal distance wandered away from a food source. Virus-exposed larvae exhibited increased maximum distance of horizontal movement compared to the control, but did not significantly differ in the probability of wandering versus no movement. We also found variation in the distance wandered among the five virus isolates. However, grouping the five isolates into two previously-described viral strains or genogroups did not improve predicted differences in movement, perhaps due to the presence of within-strain genetic variation among isolates in the viral genes involved in controlling host behavior. Further work is needed to determine whether the observed between-isolate variation is the result of adaptive evolution. These results suggest that the NPV infecting *D. vanillae* manipulates larval behavior to increase horizontal wandering, which could lead to higher pathogen fitness by increasing long-distance dispersal of the virus across the landscape.

## Introduction

An extended phenotype is the manipulation of behavioral or physical traits of a host by a parasite or pathogen, such that the host organism is used as a means of expressing the pathogen’s genes (Dawkins 1982). This manipulation will then increase the parasite’s ability to reproduce or pass to the next host, often by altering the locomotory or climbing behavior of the host (Hoover et al. 2011, Gasque et al. 2019). Sometimes referred to as “zombie behavior,” extended phenotypes have been observed in many host-parasite relationships in insects, from fungal pathogens (Andersen et al. 2009) to helminths (Biron et al. 2006) to viruses (Hoover et al. 2011). Parasite-induced behavior are considered to meet the criteria for an extended phenotype when a behavior 1) was due to the manipulation of parasites 2) increased parasite fitness (i.e., transmission) and 3) decreased host fitness (Andersen et al. 2009).

Another common example of an extended phenotype occurs in nucleopolyhedrosis virus (NPV), a baculovirus which infects Lepidopteran larvae (Harrison and Hoover 2012, Williams et al. 2017, Gasque et al. 2019). A larva becomes infected after ingesting virus-contaminated leaf material. Following infection with NPV, the host exhibits abnormal behavior, including climbing to elevated positions (“tree-top disease” or *Wipfelkrankheit*; Hofmann 1891), a phenomenon widely associated with baculovirus infection (Goulson 1997, Ros et al. 2015, Gasque et al. 2019), and thought to be due to an induced phototropic response in infected larvae (van Houte, van Oers, et al. 2014). At the end of the infectious cycle, the host dies and liquifies, releasing millions of viral particles into the environment. Viral particles are polyhedral occlusion bodies (OBs), comprising a protective protein envelope containing virions, which allows for the baculovirus to persist in the environment outside the host (Rohrmann 2019). The virus-induced elevated positions at death are thought to increase transmission, as virus from the liquified host drips down to cover a larger area of leaf material where it can be consumed by a new host (Goulson et al. 1995). Exposed *Mamestra brassicae* larvae (Lepidoptera: Noctuidae) were also more likely to position themselves on the upper side of leaves when infected, further increasing the spread of virus (Goulson 1997). In another Noctuid moth, *Spodoptera exigua*, the longer a larva remained infected with NPV, the higher it climbed (van Houte et al. 2014), as expected for a behavior that is induced by infection. In addition to the well-documented vertical climbing behavior caused by NPV infection, there is also evidence of NPV-induced horizontal movement, also called “enhanced locomotory activity” (Kamita et al. 2005, Katsuma et al. 2012, van Houte et al. 2012). Similar to inducing vertical movement, increasing horizontal wandering may benefit virus fitness by increasing dispersal, allowing the virus to exploit new hosts over a larger geographic area. Horizontal movements could also help sustain the viral populations over time, particularly in environments where the host population fluctuates (Berryman 1996).

The viral-induced behavioral changes suggest an underlying genetic basis for host manipulation in NPVs. The viral gene *egt* (ecdysteroid UDP-glucosyltransferase gene), which encodes an enzyme that inactivates the larval molting hormone 20-hydroxyecdysone, induces altered behaviors in infected larvae (Hoover et al. 2011, Ros et al. 2015). First, exposed larvae continue to eat and grow past the point at which they would normally pause food intake for molting or pupation, leading to larger size at death, prolonged lifespan after exposure, and ultimately increased viral yield as measured by occlusion body production (O’ Reilly 1997, Ros et al. 2015). Second, *egt* is thought to be essential for the virus-induced climbing behavior in many NPV species including LdMNPV, SeMNPV, and HearNPV, with larvae exposed to an *egt*-knockout strain of NPV failing to exhibit the climbing demonstrated by those exposed to a wildtype or an *egt*-recovery strain (Hoover et al. 2011, Han et al. 2015, Wang and Hu 2019; but see Ros et al. 2015). Similar to *egt*, the viral gene *ptp* (protein tyrosine phosphatase) influences host behavior, but through an independent mechanism that enhances general locomotory activity (Kamita et al. 2005, van Houte et al. 2012, van Houte, Ros, et al. 2014). This gene, thought to be originally derived from Lepidopteran hosts, is present in all group 1 and some group 2 Alphabaculoviurses, with two copies present in some group 1 species (*ptp-1* and *ptp-2*; van Houte et al. 2012). The *ptp* gene in baculoviruses co-opts the natural host wandering behavior that occurs prior to molt, by triggering horizontal movement in infected larvae 12-24 hours prior to virus-induced death (Kamita et al. 2005). The increased horizontal movement increases the area contaminated by viral particles upon larval death, further increasing viral transmission. Additionally, exposed larvae are thought to leak virus particles as they move, further spreading the virus (van Houte et al. 2012).

While a genetic basis for virus-induced movement behavior is well-established in NPVs, no studies have yet compared movement among naturally occurring isolates. NPVs are highly genetically variable, with multiple isolates coexisting within a single host population (Fleming-Davies and Dwyer 2015), across several geographic regions (Escribano et al. 1999, Cory et al. 2005), or even within a single exposed individual (Erlandson 2009). Baculoviruses are known to undergo frequent recombination events, particularly among closely-related strains, leading to high genetic diversity (Federici 1997). Furthermore, genetically distinct strains often show phenotypic differences, including speed of kill and transmission rates (Escribano et al. 1999, Cory et al. 2005, Fleming-Davies and Dwyer 2015, Hudson et al. 2016).

We studied whether naturally occurring isolates of the NPV that infects *Dione (Agraulis) vanillae* (Nymphalidae) larvae vary in the wandering behavior they induce in infected larvae. The nucleopolyhedrosis virus (NPV) that infects *D. vanillae* (DijuNPV, also called AgvaNPV) is an Alphabaculovirus which infects *D. vanillae* as well as several related Lepidopteran species (Nymphalidae) in its native range (Rodríguez et al. 2011, Ribeiro et al. 2019). This NPV exhibits extensive genetic variability within the free living host population of *D. vanillae* in the County of San Diego, where this study was conducted. Within San Diego County, we have observed two main strains or clades of the virus, which vary phenotypically between strains in their infection rates as well as genetically in key life history genes including behavior-controlling genes *egt* and *ptp-1* (Kokusho and Katsuma 2021). While one of the strains (North County) is relatively homogeneous genetically among isolates, the other strain (City of San Diego) has substantial variation among isolates within the clade. In the current study, we used laboratory infections with five different field-collected isolates across two strains to quantify variation in horizontal movements of infected *D. vanillae* larvae.

## Methods

### Study system and insect rearing

The Gulf Fritillary butterfly, *Dione (Agrualis) vanillae* (Nymphalidae) is a butterfly species native to South and Central America, the West Indies and into the Southern United States. Larvae rely on *Passiflora* species as their food plants (Daniels 2009). *D. vanillae* has been established in San Diego, CA for several hundred years, in free living butterfly populations that depend on cultivated *Passiflora spp.* plants for food (Halsch et al. 2020). To obtain the virus isolates used, larvae were collected from natural populations at several field locations across San Diego and raised to adulthood. Any larva that died of virus was collected, and occlusion bodies were isolated to be used for infection (Rodríguez et al. 2011). Isolates used in this experiment were previously subject to whole genome sequencing in another study and thus were confirmed to be genetically distinct (see *Laboratory infections* below for details).

Larvae for use in experimental infections were first generation offspring of field-collected *D. vanillae*, bred and reared in the lab to ensure that they had not previously been exposed to virus. Butterflies (n=10) were collected as larvae from several field locations across San Diego and allowed to mate freely in a single population. Larvae were transferred at the first instar from the mating cage to metal cages surface and raised in groups of 25 on surface sterilized *Passiflora caerulea* leaf material. Ten percent household bleach solution (0.4% sodium hypochlorite) was used for surface sterilization. Butterflies and larvae were housed in environmental chambers at either ambient temperature (25° C) or high temperature (35° C) with 70% humidity on a 12 hour light/dark cycle prior to infection (see below). Environmental chambers were maintained with the described temperature, photoperiod and humidity settings for the duration of the experiment, including during the maze assays after infection. We found no effect of rearing temperature on wandering behavior (see Supplement), and thus have analyzed all data for both temperature treatments pooled (25° C and 35° C).

### Laboratory infections

All infections were conducted using fourth instar caterpillars that molted in the last 24 hours. The larvae go through five instars before pupation, and fourth instar was chosen for infection because larvae are less susceptible to the virus compared to smaller instars, allowing them to survive longer post exposure (typically 5-10 days in other NPVs, Engelhard and Volkman 1995, Federici 1997) and thus giving more time for observations of wandering behavior. Larvae were infected in the laboratory using a modified droplet feeding method on leaf discs (Elderd et al. 2013). Doses of virus solution containing 200 or 1000 occlusion bodies suspended in Milli-Q water (higher dose used in year 1 experiments only; n=22 larvae) were prepared using stock virus solutions of field collected isolates in water (e.g., 200 OBs dose is equivalent to 6.7x10^4^ OBs/mL concentration). Haemocytometer counts (n=12 counts per sample, averaged) were used to determine concentration of occlusion bodies in stock solution. To infect larvae, a water control or three uL of virus solution was pipetted onto 6mm *Passiflora caerulea* leaf disks, cut using a 10% bleach sterilized leather punch. A single caterpillar was placed in a 2oz plastic cup containing an infected or control leaf disk depending on treatment group. The larval cups were placed in environmental chambers (see *Study system and insect rearing* above) on damp paper towels to prevent drying of the leaf disks. Caterpillars were given up to 24 hours to consume the entire leaf disk; individuals that had not consumed the entire leaf disk in this period were excluded from the experiment. Once the discs were consumed, individual larvae were transferred to mazes (one larva per maze; see details below) until pupation or death from virus. Control larvae that showed signs of infection or did not successfully emerge as adults were removed from the analysis.

The experiment took place over two years with a total sample size of 175 larvae divided into 5 isolates: OLE-159, BFF-004, CDO-153, GRM-060, TRV-277. These isolates are categorized into two genetically and phenotypically distinct strains based on prior work by one of the authors (Fleming-Davies et al. in prep) that found that viral whole genomes clustered into two distinct clades with different transmission rates: North County (OLE-159, BFF-004), with a lower mean but less variable infection rate and San Diego County (CDO-153, GRM-060, TRV-277), with a higher mean infection but more variable transmission. The five isolates used in this study were chosen to represent both major strains as well as between- and within-strain variation in the *egt* and *ptp* genes (Figure 1), the two viral genes that have been previously documented to control larval behavior in related NPVs (Hoover et al. 2011, van Houte et al. 2012).

**Figure 1:**
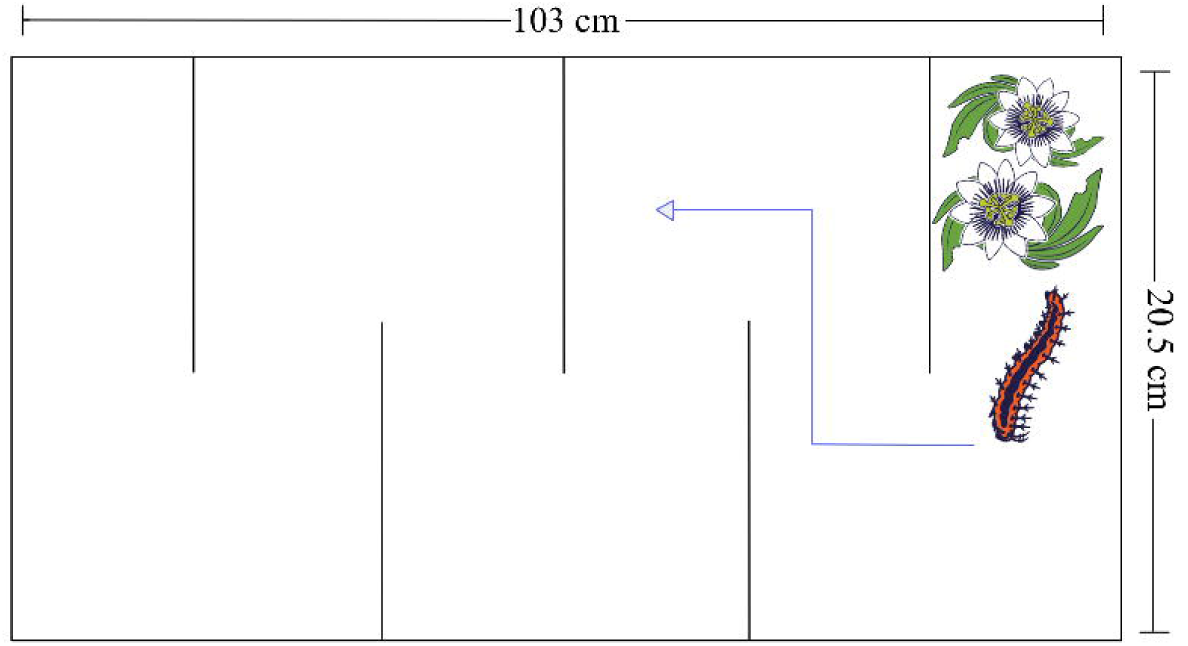
Heatmap representing differences in the nucleotide sequence of the a) *egt* gene and b) *ptp* gene among experimental isolates of *AgvaNPV* used in this study. The reference genome shown here is from isolates collected in *Dione juno* (Ribeiro et al. 2019). Sequence differences between isolates were inferred using a multiple sequence alignment with MAFFT in Geneious Prime 2026.0.2. (Katoh and Standley 2013).

### Maze Assay for Horizontal Movement

To measure larval horizontal movement, we built twenty-five 12.5 cm tall by 103 cm long mazes using acrylic plexiglass, with five walls creating the maze path (Figure 2). Mazes were assembled with autoclavable metal rods, and thus mazes were reused between larvae after disassembly, surface sterilization with 10% household bleach solution, and reassembly. A single larva was placed into the right side of each maze, containing larval food consisting of a virus-free vine of *Passiflora caerulea*. Thus, any movement out of this “home” area is more likely to be due to the virus, as larvae should not choose to leave the only food source until preparing for pupation. The position of the larvae were measured daily, until they died of virus or pupated. Each maze was kept in an environmental chamber at ambient (25° C) or elevated (35° C) temperature, 70% humidity and a 12 hour light/dark cycle as described above until the larva died of virus exposure or pupated.

**Figure 2:**
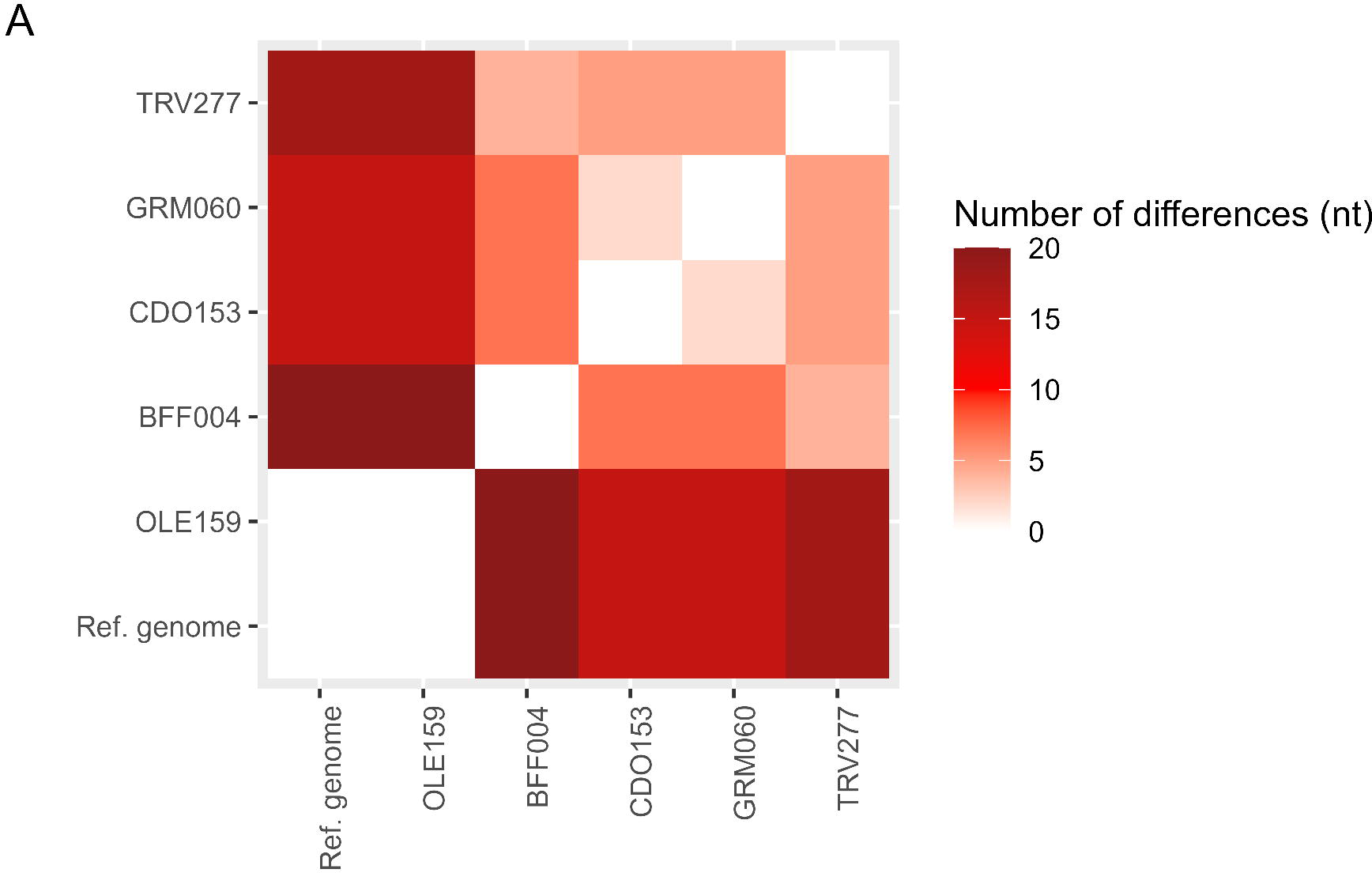
Diagram of the plexiglass maze designed to test horizontal movements. Larvae were introduced into the maze on the leaf material contained in the far right area, and wandering was measured as distance moved beyond that area. Larva and plant drawings by Kieran Gomez.

Infection and strain effects on the maximum distance wandered were analyzed using a Generalized Linear Model (GLM) with a zero-inflated Gaussian distribution in R (glmmTMB package in R; (Brooks et al. 2017)). This distribution was chosen due to the excess number of non-wandering individuals observed in the dataset, as well as to appropriately model the non-zero values as continuous data following a normal distribution (maximum distance was log transformed to meet assumptions of normality). Specifically, we examined 1) differences between exposed vs unexposed control individuals, 2) differences among the five virus isolates, and 3) differences between the two virus strains, each containing multiple isolates. We also considered additional correlates of viral dose, rearing temperature, and fate of virus-exposed larvae (i.e., successful pupation or virus-induced larval death).

## Results

Virus-exposed larvae wandered significantly farther than controls (Figure 3; GLM with zero-inflated Gaussian distribution of log transformed horizontal distance traveled: Effect of virus exposure: 3.25, LR=16.7, df=1, p<0.001). Furthermore, the horizontal distance moved varied between the different virus isolates (Fig 3; GLM with zero-inflated Gaussian distribution of log transformed distance; isolate effect compared to exposed-only model: LR=10.34, df=4, p=0.035), although the strain effect was marginally nonsignificant, due to variation among isolates within the City of San Diego strain (strain effect compared to exposed-only model: LR=3.71, df=1, p=0.05). However, virus exposure did not significantly increase the probability of any horizontal wandering versus no movement (25 % of exposed individuals versus 10% of control individuals wandered; Fisher’s exact test for exposed larvae versus unexposed controls, odds ratio=2.92, p=0.23). Similarly, there was no difference in the probability of wandering versus no movement among different strains (Figure 4; Fisher’s exact test, p=0.34; odds ratio not applicable for > 2x2 matrix) or isolates (Fisher’s exact test, p=0.64). Across all isolates, 64 out of 77 larvae (0.83; 95% binomial confidence intervals: 0.73, 0.90) of the larvae infected died of virus (see Table S1). Fourteen infected insects survived until pupation, but only 9 of these emerged successfully as butterflies. Eventual fate of exposed larvae (virus-killed or pupated) did not predict wandering behavior (LR=0.989, df=1, p= 0.32). Viral dose and rearing temperature also had no significant effect on distance wandered (dose: LR=0.937, df=1, p= 0.33; temperature: LR=2.05, df=1, p=0.15).

**Figure 3:**
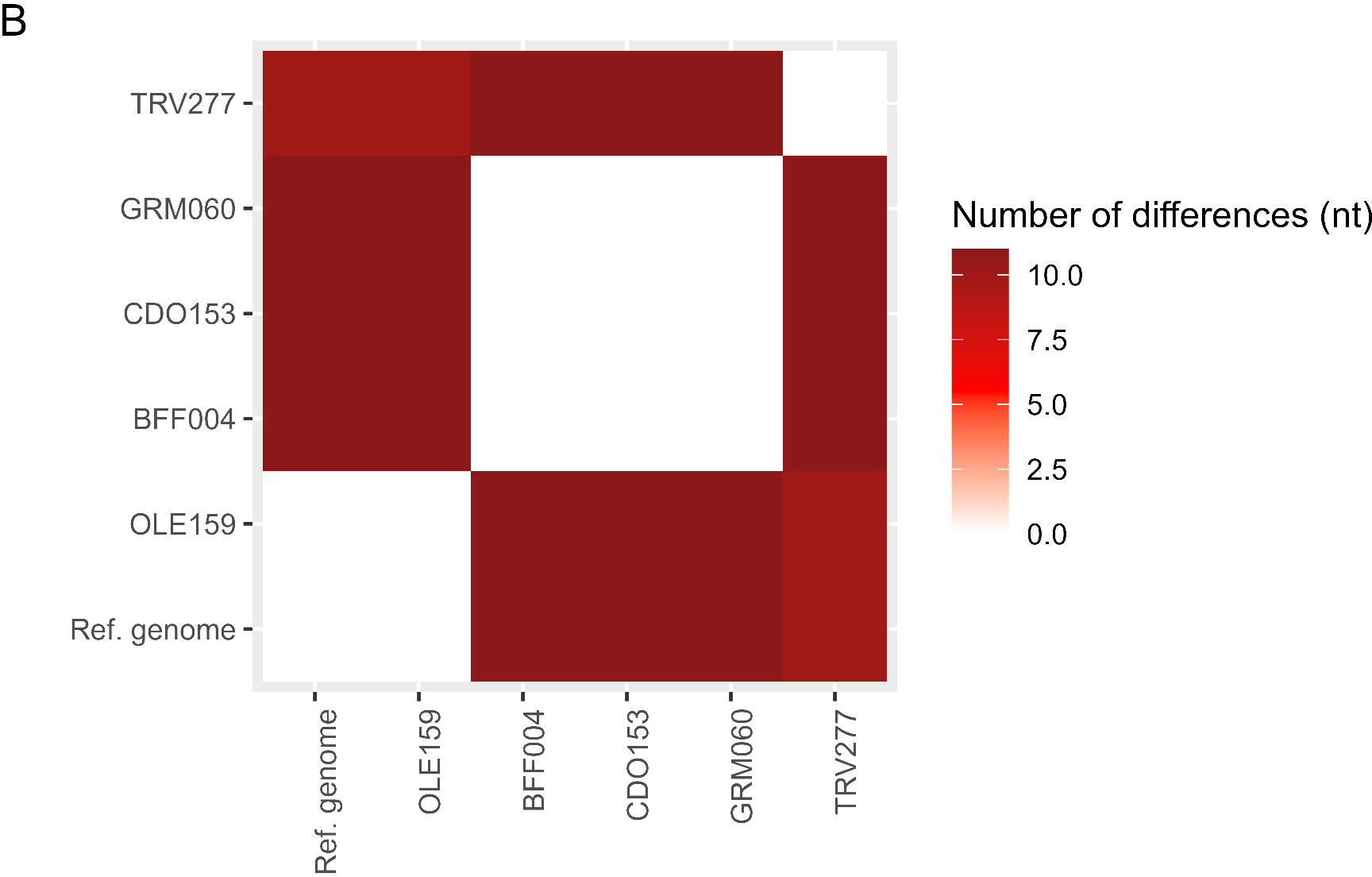
Strip chart of maximum horizontal distance for each larva exposed to one of five virus isolates (OLE-159: n=11, BFF-004: n=19, CDO-153:n=14, GRM-060: n=19, TRV-277: n=24 larvae) or unexposed controls (n=20 larvae).

**Figure 4:**
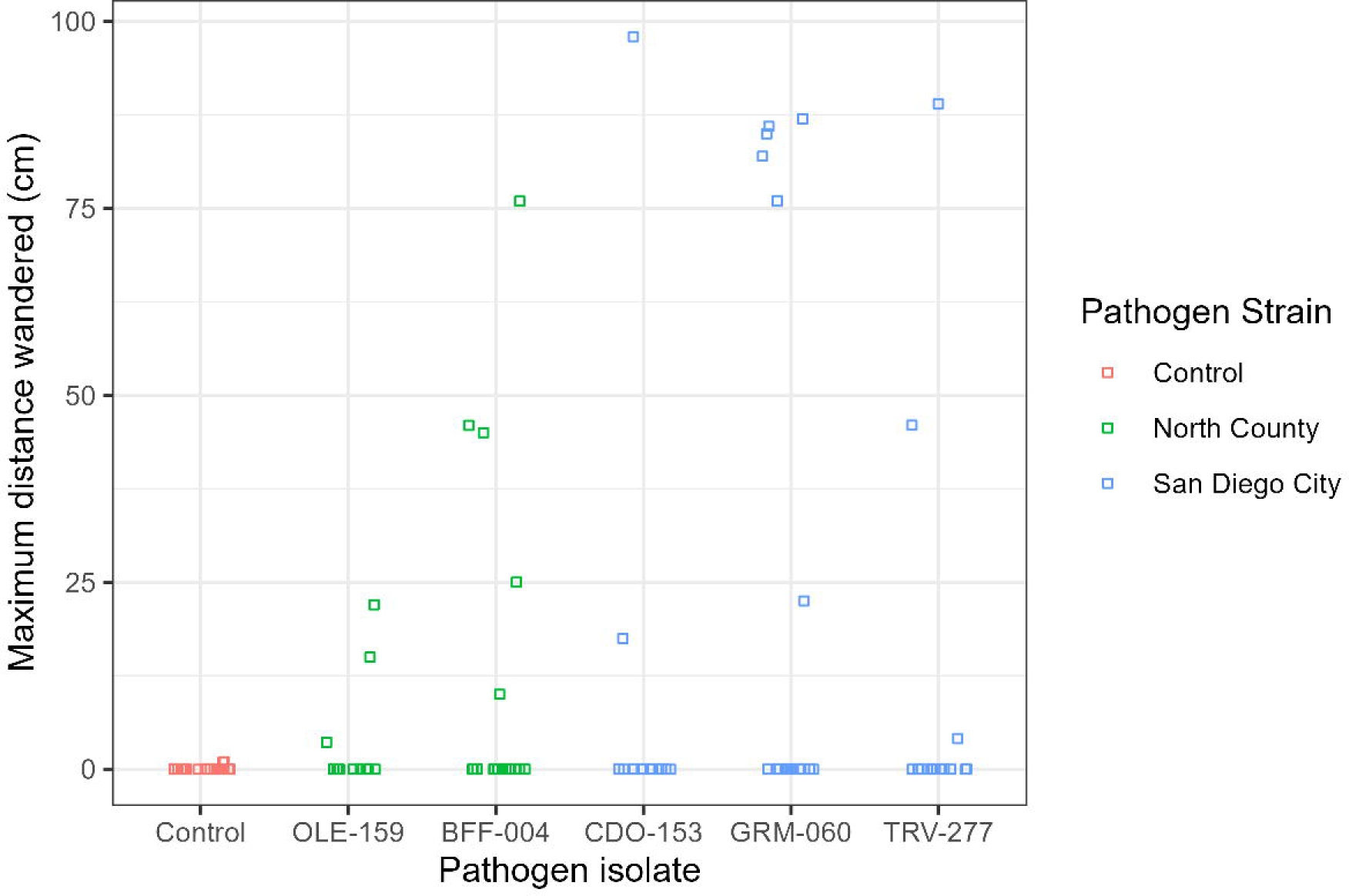
Mosaic plot of larval wandering (movement beyond food source) for unexposed control larvae (n=20) compared to those exposed to the North County (n=30) or San Diego City (n=48) NPV strains.

## Discussion

Our finding that exposed larvae traveled farther horizontally than unexposed control larvae suggests that the NPV that infects *D. vanillae* larvae may stimulate horizontal movements to increase viral transmission as seen in other NPVs of Lepidoptera (Kamita et al. 2005, Katsuma et al. 2012, van Houte et al. 2014). We also found significant differences in the distance traveled among larvae exposed to different virus isolates collected within San Diego County. Notably, the City of San Diego isolates (GRM-060, CDO-153, and TRV-277) that showed variation in the distance traveled also showed variation in their *egt* gene sequences, whereas the North County isolates (BFF-004 and OLE-159) that showed no *egt* variation were similar in their distances traveled. GRM-060, the isolate with the longest distances traveled, also had the most differences in its *egt* sequence from the other isolates (Figure 1A), which could indicate that certain *egt* variants directly affect horizontal movement as well as vertical movement. While the main focus of this paper was on horizontal movement, vertical movement was measured for a subset of individuals (n=57; Figure S1), and was positively but not significantly correlated with horizontal distance traveled (Spearman’s correlation between vertical and horizontal distance traveled: 0.483, bootstrapped 95% CI: -0.25, 0.89). Since the focus of our study was measuring horizontal movement, the height of the maze was only 12.5 cm, and many of the individuals that moved horizontally were also found near the maximum height, suggesting that the height of the enclosure may have limited our ability to measure virus-induced climbing. Further evidence, ideally through manipulation of *egt* gene sequences on an otherwise common viral genetic background (similar to the work of (Hoover et al. 2011)), would help determine whether *egt* alone or in combination with other genes may explain this behavioral variation.

Differences in mean infection rates between the viral strains might also drive part or all of the observed variation in wandering behavior across isolates: larvae were exposed to a constant 200 OB dose, but this is effectively a ‘stronger’ dose (i.e., resulting in higher infection rates; Table S1) for the isolates in the City of San Diego group, and thus could contribute to the greater distances moved, for example if the viral load within the host was maintained at higher levels throughout the course of infection. It is also possible that another viral gene that directly manipulates behavior, potentially one not yet identified, could be influencing movement.

Interestingly, while *egt* sequences varied among City of San Diego isolates, their *ptp-1* sequences remained highly similar (Figure 1B), further suggesting *egt* or an uncharacterized gene as a likely driver of this behavioral difference. Although the roles of *egt* and *ptp* in controlling host behavior are well-documented in the literature, other work has measured increased vertical and horizontal movement in larvae infected with NPV isolates without functional *egt* or *ptp* genes in *Bombyx mori* nucleopolyhedrovirus (Kokusho and Katsuma 2021), suggesting the potential for alternative undescribed genetic mechanisms for inducing these behaviors.

Wandering, as measured in our study, was a rare behavior (21 of 97 individuals, or 22% of all larvae, Figure 4), despite high infection rates in exposed larvae (82% across all isolates, minimum 71% in the least-virulent isolate; Table S1). While larvae exposed to the virus appeared to move more than unexposed controls, this trend was not statistically significant, suggesting that any effect of virus exposure on the choice to wander or not may vary with other factors or require greater sample size to detect. Photoperiod, directionality of light exposure, and time since infection have all been shown to be important in inducing climbing and/or wandering behavior (van Houte, van Oers, et al. 2014, Han et al. 2018, Liu et al. 2022). We maintained a constant 12-hour photoperiod and light from directly above the mazes in the environmental chambers to control for any effect of light exposure. It is possible that manipulating the angle of exposure (i.e., to mimic variation of sun angle with time of day) or orienting the mazes with the light source away from the larval food would increase the prevalence of virus-induced movement.

Insights into NPV control of host behavior could also aid in the development of more effective pest management. Baculoviruses are potential insect control agents, as they are able to reduce the impact of pest-related damage while offering advantages over chemical pesticides: they are natural pathogens already prevalent in the environment, typically highly host-specific and thus harmless to non-target species, and capable of persisting in the environment for several years (Moscardi 1989, Miller 1997). However, naturally occurring baculoviruses tend to have a slower speed of kill compared to chemical pesticides (Beas-Catena et al. 2014). Understanding the effects of genetic variation in key viral genes such as egt and ptp is essential in developing baculovirus-based pesticide (Black et al. 1997, Emara el-Wakeil et al. 2020). For example, *egt* is often deleted from commercial baculovirus strains because this reduces host feeding significantly and results in a shorter lifespan,both of which are beneficial for biocontrol of agricultural pests (Beas-Catena et al. 2014). However, *egt*-deletion comes with multiple downsides: eliminating virus-induced host movement and reducing occlusion body yields, which might reduce natural spread of the NPV, thus requiring more frequent application and leading to increased economic cost to farmers.

Another potential improvement to biocontrol with NPVs is the use of multiple strains together (Páez and Fleming-Davies 2020). However, for this to be effective, the strains used must be phenotypically different, such as having distinct transmission rates or infection variability. Phenotypically distinct strains are predicted to complement each other, ultimately lowering the host population beyond either strain alone (Páez and Fleming-Davies 2020). More research is needed to fully understand these viral-host interactions and their implications for pest management and disease control. For example, exploring the genetic basis of phenotypic differences, such as how the behavior induced by *egt* and *ptp* genes influence transmission rates, could help identify optimal strains combinations to use as biocontrol agents.

Finally, the interaction between baculoviruses and their hosts play a pivotal role in natural ecological dynamics such as population cycles of outbreaking insects (Cooper et al. 2003, Hwang et al. 2023). The changes in behavior induced by the virus, such as the “tree-top” phenomenon and the increased horizontal movements, could influence these population cycles and landscape-scale movements of hosts and pathogens, particularly if we consider the three dimensional space used. This is particularly relevant for viruses infecting insect host species that use tall trees as larval food plants; varied pathogen strategies for manipulation of host movement could potentially allow for vertical stratification of hosts by different virus strains, perhaps increasing the chance of coexistence by multiple competing pathogen strains. Future work is needed to examine whether among-strain variation in pathogen strategies for host behavioral manipulation could thus be an additional mechanism maintaining pathogen variation within host populations.

## Supporting information

Supplemental results

## Acknowledgments

Many thanks to Alexa Alderete, Felix Turrubiartes, Kailani Topasna, Allison Dela Cruz, Damian Osuna, Brianna Leveille, Ethan Huynh, Eclas Adde, Jocelyne Castro Luquin, and Alyssa Sanchez for help conducting laboratory experiments, and to Steven Saxer, Mechanical Shop Manager at USD Shiley-Marcus School of Engineering for help with maze design and construction.

---

**Figure S1:**
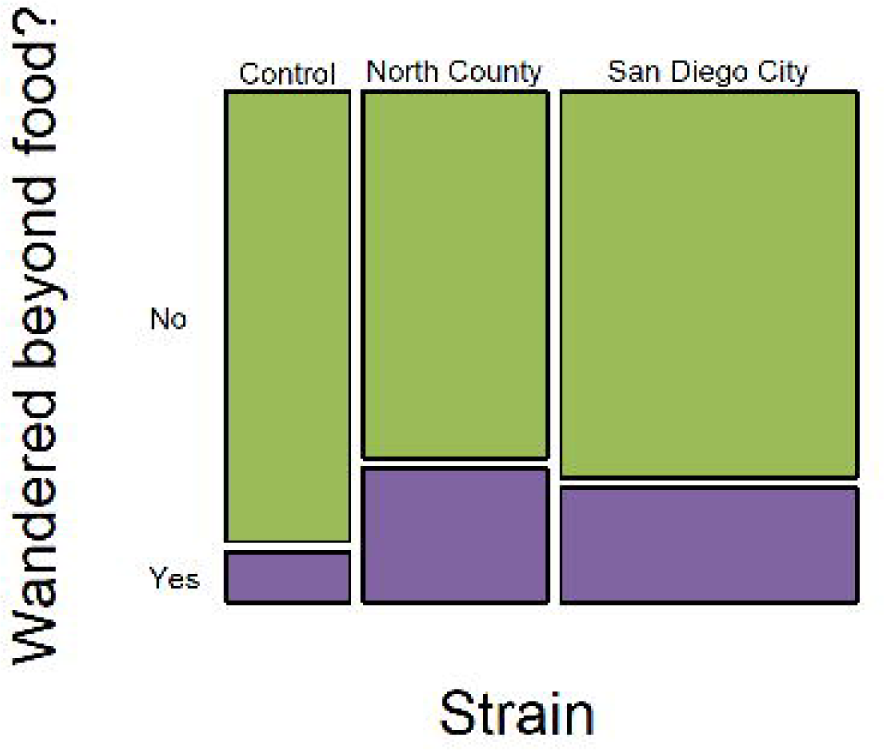
Maximum height versus maximum distance wandered for a subset of individuals with vertical height measurements. Points show only those larvae with wandering distance greater than 0 and measured height (n=9). Height of the enclosure was 12.5 cm.

## References

Andersen SB, Gerritsma S, Yusah KM, et al. 2009. The Life of a Dead Ant: The Expression of an Adaptive Extended Phenotype. Am. Nat. 174(3):424–433. 10.1086/603640

Beas-Catena A, Sánchez-Mirón A, García-Camacho F, et al. 2014. BACULOVIRUS BIOPESTICIDES: AN OVERVIEW. J. Anim. Plant Sci. 24(2):362–373.

Berryman AA. 1996. What causes population cycles of forest Lepidoptera? Trends Ecol. Evol. 11(1):28–32. 10.1016/0169-5347(96)81066-4

Biron DG, Ponton F, Marché L, et al. 2006. ‘Suicide’ of crickets harbouring hairworms: a proteomics investigation. Insect Mol. Biol. 15(6):731–742. 10.1111/j.1365-2583.2006.00671.x

Black BC, Brennan LA, Dierks PM, et al. 1997. Commercialization of Baculoviral Insecticides. In: Miller LK, editor. The Baculoviruses. Boston, MA: Springer US. p. 341–381. 10.1007/978-1-4899-1834-5

Brooks M E, Kristensen K, Benthem K J, van, et al. 2017. glmmTMB Balances Speed and Flexibility Among Packages for Zero-inflated Generalized Linear Mixed Modeling. R J. 9(2):378. 10.32614/RJ-2017-066

Cooper D, Cory JS, Theilmann DA, et al. 2003. Nucleopolyhedroviruses of forest and western tent caterpillars: cross-infectivity and evidence for activation of latent virus in high-density field populations. Ecol. Entomol. 28(1):41–50. 10.1046/j.1365-2311.2003.00474.x

Cory JS, Green BM, Paul RK, et al. 2005. Genotypic and phenotypic diversity of a baculovirus population within an individual insect host. J. Invertebr. Pathol. 89(2):101–111. 10.1016/j.jip.2005.03.008

Daniels JC. 2009. Gulf Fritillary Butterfly, Agraulis vanillae (Linnaeus) (Insecta: Lepidoptera: Nymphalidae): EENY 423/IN804, 2/2009. EDIS. 2009(2). 10.32473/edis-in804-2009

Dawkins R. 1982. The extended phenotype: the gene as the unit of selection. Oxford [Oxfordshire]; San Francisco: Freeman.

Elderd BD, Rehill BJ, Haynes KJ, et al. 2013. Induced plant defenses, host–pathogen interactions, and forest insect outbreaks. Proc. Natl. Acad. Sci. 110(37):14978–14983. 10.1073/pnas.1300759110

Emara el-Wakeil NM, Saleh M, Abu-hashim M, editors. 2020. Cottage Industry of Biocontrol Agents and Their Applications: Practical Aspects to Deal Biologically with Pests and Stresses Facing Strategic Crops. 1st ed. 2020. Cham: Springer (Springer eBooks Earth and Environmental Science).

Engelhard EK, Volkman LE. 1995. Developmental Resistance in Fourth Instar Trichoplusia ni Orally Inoculated with Autographa californica M Nuclear Polyhedrosis Virus. Virology. 209(2):384–389. 10.1006/viro.1995.1270

Erlandson MA. 2009. Genetic variation in field populations of baculoviruses: Mechanisms for generating variation and its potential role in baculovirus epizootiology. Virol. Sin. 24(5):458–469. 10.1007/s12250-009-3052-1

Escribano A, Williams T, Goulson D, et al. 1999. Selection of a Nucleopolyhedrovirus for Control of Spodoptera frugiperda (Lepidoptera: Noctuidae): Structural, Genetic, and Biological Comparison of Four Isolates from the Americas. J. Econ. Entomol. 92(5):1079–1085. 10.1093/jee/92.5.1079

Federici BA. 1997. Baculovirus Pathogenesis. In: Miller LK, editor. The Baculoviruses. Boston, MA: Springer US. p. 33–59. 10.1007/978-1-4899-1834-5_3

Fleming-Davies AE, Dwyer G. 2015. Phenotypic Variation in Overwinter Environmental Transmission of a Baculovirus and the Cost of Virulence. Am. Nat. 186(6):797–806. 10.1086/683798

Gasque SN, Van Oers MM, Ros VI. 2019. Where the baculoviruses lead, the caterpillars follow: baculovirus-induced alterations in caterpillar behaviour. Curr. Opin. Insect Sci. 33:30–36. 10.1016/j.cois.2019.02.008

Goulson D. 1997. Wipfelkrankheit : modification of host behaviour during baculoviral infection. Oecologia. 109(2):219–228. 10.1007/s004420050076

Goulson D, Hails RS, Williams T, et al. 1995. Transmission Dynamics of a Virus in a Stage-Structured Insect Population. Ecology. 76(2):392–401. 10.2307/1941198

Halsch CA, Shapiro AM, Thorne JH, et al. 2020. A winner in the Anthropocene: changing host plant distribution explains geographical range expansion in the gulf fritillary butterfly. Ecol. Entomol. 45(3):652–662. 10.1111/een.12845

Han Y, Van Houte S, Drees G, et al. 2015. Parasitic Manipulation of Host Behaviour: Baculovirus SeMNPV EGT Facilitates Tree-Top Disease in Spodoptera exigua Larvae by Extending the Time to Death. Insects. 6(3):716–731. 10.3390/insects6030716

Han Y, Van Houte S, Van Oers MM, et al. 2018. Timely trigger of caterpillar zombie behaviour: temporal requirements for light in baculovirus-induced tree-top disease. Parasitology. 145(6):822–827. 10.1017/S0031182017001822

Harrison R, Hoover K. 2012. Baculoviruses and Other Occluded Insect Viruses. In: Insect Pathology. Elsevier. p. 73–131. 10.1016/B978-0-12-384984-7.00004-X

Hoover K, Grove M, Gardner M, et al. 2011. A Gene for an Extended Phenotype. Science. 333(6048):1401–1401. 10.1126/science.1209199

van Houte S, van Oers MM, Han Y, et al. 2014. Baculovirus infection triggers a positive phototactic response in caterpillars to induce ‘tree-top’ disease. Biol. Lett. 10(12):20140680. 10.1098/rsbl.2014.0680

van Houte S, Ros VID, Mastenbroek TG, et al. 2012. Protein Tyrosine Phosphatase-Induced Hyperactivity Is a Conserved Strategy of a Subset of BaculoViruses to Manipulate Lepidopteran Host Behavior. Breuker C, editor. PLoS ONE. 7(10):e46933. 10.1371/journal.pone.0046933

van Houte S, Ros VID, Van Oers MM. 2014. Hyperactivity and tree-top disease induced by the baculovirus AcMNPV in Spodoptera exigua larvae are governed by independent mechanisms. Naturwissenschaften. 101(4):347–350. 10.1007/s00114-014-1160-8

Hudson AI, Fleming-Davies AE, Páez DJ, et al. 2016. Genotype-by-genotype interactions between an insect and its pathogen. J. Evol. Biol. 29(12):2480–2490. 10.1111/jeb.12977

Hwang H, Acharya R, Lucas MDC, et al. 2023. Effects of *Lymantria dispar* multiple nucleopolyhedrovirus and *Bacillus thuringiensis* var. *kurstaki* on different larval instars of *Lymantria dispar asiatica*. Arch. Insect Biochem. Physiol. 113(1):e22002. 10.1002/arch.22002

Kamita SG, Nagasaka K, Chua JW, et al. 2005. A baculovirus-encoded protein tyrosine phosphatase gene induces enhanced locomotory activity in a lepidopteran host. Proc. Natl. Acad. Sci. 102(7):2584–2589. 10.1073/pnas.0409457102

Katoh K, Standley DM. 2013. MAFFT Multiple Sequence Alignment Software Version 7: Improvements in Performance and Usability. Mol. Biol. Evol. 30(4):772–780. 10.1093/molbev/mst010

Katsuma S, Koyano Y, Kang W, et al. 2012. The Baculovirus Uses a Captured Host Phosphatase to Induce Enhanced Locomotory Activity in Host Caterpillars. Schneider DS, editor. PLoS Pathog. 8(4):e1002644. 10.1371/journal.ppat.1002644

Kokusho R, Katsuma S. 2021. Bombyx mori nucleopolyhedrovirus ptp and egt genes are dispensable for triggering enhanced locomotory activity and climbing behavior in Bombyx mandarina larvae. J. Invertebr. Pathol. 183:107604. 10.1016/j.jip.2021.107604

Liu Xiaoming, Tian Z, Cai L, et al. 2022. Baculoviruses hijack the visual perception of their caterpillar hosts to induce climbing behaviour thus promoting virus dispersal. Mol. Ecol. 31(9):2752–2765. 10.1111/mec.16425

Miller LK, editor. 1997. The Baculoviruses. Boston, MA: Springer US. 10.1007/978-1-4899-1834-5

Moscardi F. 1989. Use of viruses for pest control in Brazil: the case of the nuclear polyhedrosis virus of the soybean caterpillar, Anticarsia gemmatalis. Mem. Inst. Oswaldo Cruz. 84:51–56.

O’ Reilly DR. 1997. Auxiliary Genes of Baculoviruses. In: Miller LK, editor. The Baculoviruses. Boston, MA: Springer US. p. 267–296. 10.1007/978-1-4899-1834-5

Páez DJ, Fleming-Davies AE. 2020. Understanding the Evolutionary Ecology of host–pathogen Interactions Provides Insights into the Outcomes of Insect Pest Biocontrol. Viruses. 12(2):141. 10.3390/v12020141

Ribeiro BM, Dos Santos ER, Trentin LB, et al. 2019. A Nymphalid-Infecting Group I Alphabaculovirus Isolated from the Major Passion Fruit Caterpillar Pest Dione juno juno (Lepidoptera: Nymphalidae). Viruses. 11(7):602. 10.3390/v11070602

Rodríguez VA, Belaich MN, Mengual Gómez DL, et al. 2011. Identification of nucleopolyhedrovirus that infect Nymphalid butterflies Agraulis vanillae and Dione juno. J. Invertebr. Pathol. 106(2):255–262. 10.1016/j.jip.2010.10.008

Rohrmann G. 2019. Baculovirus Molecular Biology. 4th ed. Bethesda (MD): National Center for Biotechnology Information (US). https://ir.library.oregonstate.edu/concern/defaults/7d279043k?localeen.

Ros VID, Van Houte S, Hemerik L, et al. 2015. Baculovirus-induced tree-top disease: how extended is the role of *egt* as a gene for the extended phenotype? Mol. Ecol. 24(1):249–258. 10.1111/mec.13019

Wang M, Hu Z. 2019. Cross-talking between baculoviruses and host insects towards a successful infection. Philos. Trans. R. Soc. B Biol. Sci. 374(1767):20180324. 10.1098/rstb.2018.0324

Williams T, Virto C, Murillo R, et al. 2017. Covert Infection of Insects by Baculoviruses. Front. Microbiol. 8:1337. 10.3389/fmicb.2017.01337

