## Supplemental results for "Wandering the NPV Maze: Effects of nucleopolyhedrovirus infection on the behavior of Gulf Fritillary (*Dione vanillae*) larvae"

**Supplementary Information**

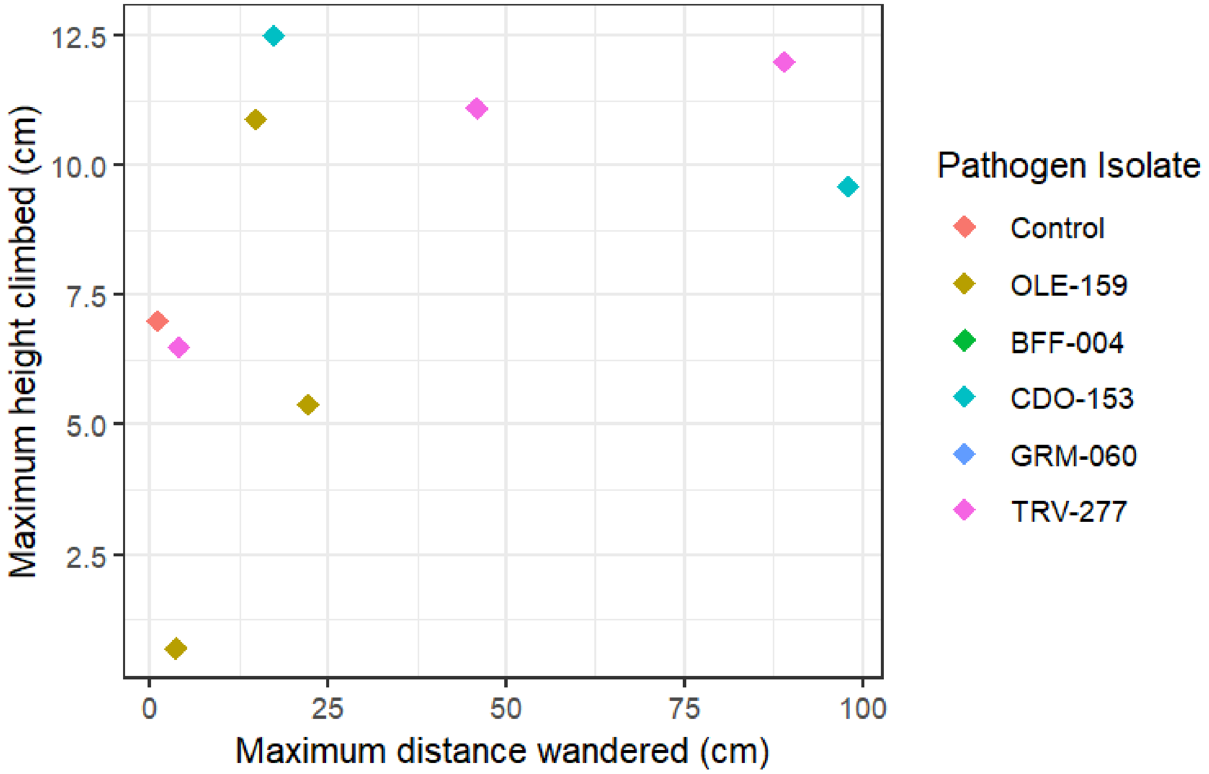


**Figure S1:**Maximum height versus maximum distance wandered for a subset of individuals with vertical height measurements. Points show only those larvae with wandering distance greater than 0 and measured height (n=9). Height of the enclosure was 12.5 cm.

**Table S1:** Summary of infection rates by isolate and dose (number of occlusion bodies, OBs). 95% binomial confidence intervals were estimated with an Agresti-Coull approximation with the package binom in R (Dorai-Raj and Cascone 2006).

| Strain | Dose (number of OBs) | Total exposed | Total infected | Proportion infected (95% confidence intervals) |
|---|---|---|---|---|
| Control | NA | 20 | 0 | 0 |
| OLE-159 | 1000 | 7 | 7 | 1.00 (0.60, 1.05) |
| | 200 | 4 | 3 | 0.75 (0.29,0.97) |
| BFF-004 | 200 | 19 | 14 | 0.74 (0.51,0.89) |
| CDO-153 | 1000 | 8 | 7 | 0.88 (0.51,1.00) |
| | 200 | 6 | 6 | 1.00 (0.56,1.05) |
| GRM-060 | 200 | 20 | 15 | 0.79 (0.56, 0.92) |
| TRV-277 | 1000 | 7 | 7 | 1.00 (0.60,1.05) |
| | 200 | 7 | 5 | 0.71 (0.35,0.92) |
| **Total exposed** | | **78** | **64** | **0.82 (0.72,0.89)** |

### References

Dorai-Raj S, Cascone AD. 2006. binom: Binomial Confidence Intervals for Several Parameterizations. :1.1-2. https://doi.org/10.32614/CRAN.package.binom
